# Long term anthropic management and associated loss of plant diversity deeply impact virome richness and composition of *Poaceae* communities

**DOI:** 10.1101/2023.01.12.523554

**Authors:** François Maclot, Virginie Debue, Carolyn Malmstrom, Denis Filloux, Philippe Roumagnac, Mathilde Eck, Lucie Tamisier, Arnaud G. Blouin, Thierry Candresse, Sébastien Massart

**Author notes:** Correspondence : Sébastien Massart.

## Abstract

Modern agriculture has influenced plant virus emergence through ecosystem simplification, introduction of new host species, and reduction in crop genetic diversity. Therefore, it is crucial to better understand virus distributions across cultivated and uncultivated communities in agro-ecological interfaces, as well as virus exchange among them. Here we advance fundamental understanding in this area by characterizing the virome of three co-occurring replicated *Poaceae* community types that represent a gradient of grass species richness and management intensity, from highly managed crop monocultures to little-managed, species-rich grasslands. We performed a large-scale study on 950 wild and cultivated *Poaceae* over two years combining untargeted virome analysis down to virus species level with targeted detection of three plant viruses. Deep sequencing revealed i) a diversified and largely unknown *Poaceae* virome (at least 51 virus species/taxa), with an abundance of so-called persistent viruses; ii) an increase of virome richness with grass species richness within the community; iii) a stable virome richness over time but a large viral intraspecific variability; and iv) contrasted patterns of virus prevalence, co-infections and geographical distribution among plant communities and species. Our findings highlight the complex structure of plant virus communities in nature and suggest the influence of anthropic management on viral distribution and prevalence.

## Introduction

Viruses are among the smallest but potentially most powerful biological entities, and significantly influence the functioning of plant communities (1). While most frequently described as pathogens, viruses may alternatively develop commensal or even mutualistic relationships with hosts (2, 3), with the nature of these relationships dependent on environmental conditions, as well as on virus and host genotypes (4, 5). In nature, plant viruses have co-evolved over long periods with their vector(s) and host(s) in complex trophic interactions networks (6). However, development of agriculture deeply modified plant communities worldwide, creating agro-ecosystems composed of both cultivated and uncultivated areas. This profound ecosystem change most certainly altered the dynamics of virus–plant–vector interactions, and likely fostered virus emergence and even disease epidemics (7, 8). To develop better mechanistic understanding of these processes requires fundamental knowledge of virus communities and their responses to host and management shifts that result from increased agricultural intensity. At present, virus dynamics are better understood within crop systems than in less managed systems, and characterization of viruses infecting wild plants is only just beginning. However, evidence indicates that virus infections are common in nature, frequently asymptomatic, and oftentimes involved multiple virus species (mixed infections) (9–11). Moreover, there is developing understanding that host diversity can powerfully influence pathogen prevalence or diversity, either increasing (amplification effect) or decreasing (dilution effect) infectious disease risk (12). Here we advance fundamental understanding of plant virus dynamics by characterizing the virome of three co-occurring *Poaceae* (or grasses) community types that represent a gradient of grass species richness and management intensity, from highly managed crop monocultures to little-managed, species-rich grasslands. The *Poaceae* virome is one of the most important to understand globally because collectively humanity depends on *Poaceae* species, directly or indirectly, for two-thirds of its caloric consumption, making this plant family essential for world food security (13, 14). Cereal crops are susceptible to at least 74 virus species worldwide (15), yield losses reaching up to 80% (16). Crop-infecting viruses may further spill into wild (non-cultivated) grasses and reduce wild host fitness, thus posing conservation risks (17–19). Alternatively, wild or weedy grasses might serve as a reservoir for crop viruses between cropping cycles. In contrast with crop viruses, only a few studies have begun to evaluate the nature and ecology of non-crop-infecting (‘wild’) viruses harbored by grasses (20, 21).

High-throughput sequencing (HTS) is well-suited for characterizing plant viruses (22). Whether plants are individually sampled and barcoded (Ecogenomics, (23)), pooled (24), or considered within spatially explicit context (Geometagenomics, (20)), viral metagenomics provides key information that can illuminate how anthropogenic perturbations impact plant-virus interactions and emergence of viral diseases (25). Here we used high-throughput sequencing (HTS) to survey, without *a priori* information, all of the viruses inhabiting the target grass communities, *i*.*e*. to characterize their viral metagenomes or viromes (26). Our first objective was to characterize the *Poaceae* viromes of our study sites, including both known and novel RNA and DNA viruses. In this effort, we strove to determine viral taxa to the finest level (i.e. species) where possible. Second, we aimed to evaluate the plant-virus richness relationship as evidenced among these plant communities and its temporal stability over two years. Third, we investigated whether the species composition of the plant community viromes was more strongly associated with community type (crop, pasture, grassland) or site location. Finally, we focused on three individual viruses to determine their prevalence and spatial distributions within plant communities, and on the haplotype level to study the genetic diversity of one latent virus within and between communities. We expected to find high virus prevalence and more severe symptoms in crop fields, while low virus prevalence associated with fewer or no visible symptoms and high co-infection levels are expected in the natural reservoir. Combining different analysis levels (plant communities, individual plants, virus species and even haplotypes), we presented a broader view of plant-virus interactions at agro-ecological landscapes. This is of particular interest as, so far, host-parasite richness relationship has been analyzed only targeting a limited number of plant viruses/hosts (27, 28) and never for the complete virome of a host community. In addition, viral metagenomics studies limited the virus diversity examination to virus family level (20) or used theoretical OTUs as an acceptable proxy to viral species (29, 30).

## Results

### A diversified and largely unknown virome in Poaceae

We characterized the virome of three co-occurring *Poaceae-*dominated communities that represent a gradient of plant species richness and management intensity (cereal crops, grazed pastures and mowed grasslands). We studied these community types across two years (2017 and 2018) at three replicated sites ten kilometers apart (Antheit, Héron, Latinne) within the Belgian National Park “Burdinale-Mehaigne” (**Figure 1**). To assess the virome in each community, we sampled 950 plants belonging to twenty-four *Poaceae* species (**Table S1**) and we used a viral metagenomic approach (virion-associated nucleic acids or VANA) on pools of 50 samples per community (**Table S2**). Librairies contained in average 17.9% of viral reads (ranging from 0 to 78%, see **Table S3**). A low level of cross-samples contamination was observed as the alien target sequences (PLRV, BSV) detected in the *Poaceae* samples represented on average 0.01% of the total reads in the libraries, which was considered as the detection threshold for the present study, meaning that any viral detection below this level was not considered. Fifty-one consensus plant virus genomes were assembled (1,496 to 13,876 nt length), covering for 47 of them all known ORFs described in the corresponding genera or families and allowing to assign them down to species level. We also detected four uncomplete virus genomes (1,497 nt to 6,785 nt length) that represented putative novel species belonging to *Closterovirus, Rymovirus, Varicosavirus* and *Amalgavirus* genera (based on ICTV demarcation criteria on RdRp/coat protein genes and host range). These fifty-one RNA viruses were assigned to sixteen virus families representing twenty-one virus genera (**Table 1**). These results demonstrate that the *Poaceae* virome remains largely unknown, with more than two thirds of the detected viruses (n = 37) corresponding to putative novel virus species (see **Table 1** and **Table S4**), in particular when it comes to persistent and mycoviruses (*i*.*e*. 3 novel alphachrysoviruses, 10 novel partitiviruses and 13 novel totiviruses), which were found in all three community types. Virome comparison from year to year revealed that 77% of the virus species were detected in both sampling years (74% for the plant viruses and 79% for the mycoviruses). This number increases to 85% when analyzing only long-term plant communities (pastures and grasslands).

**Table 1.**
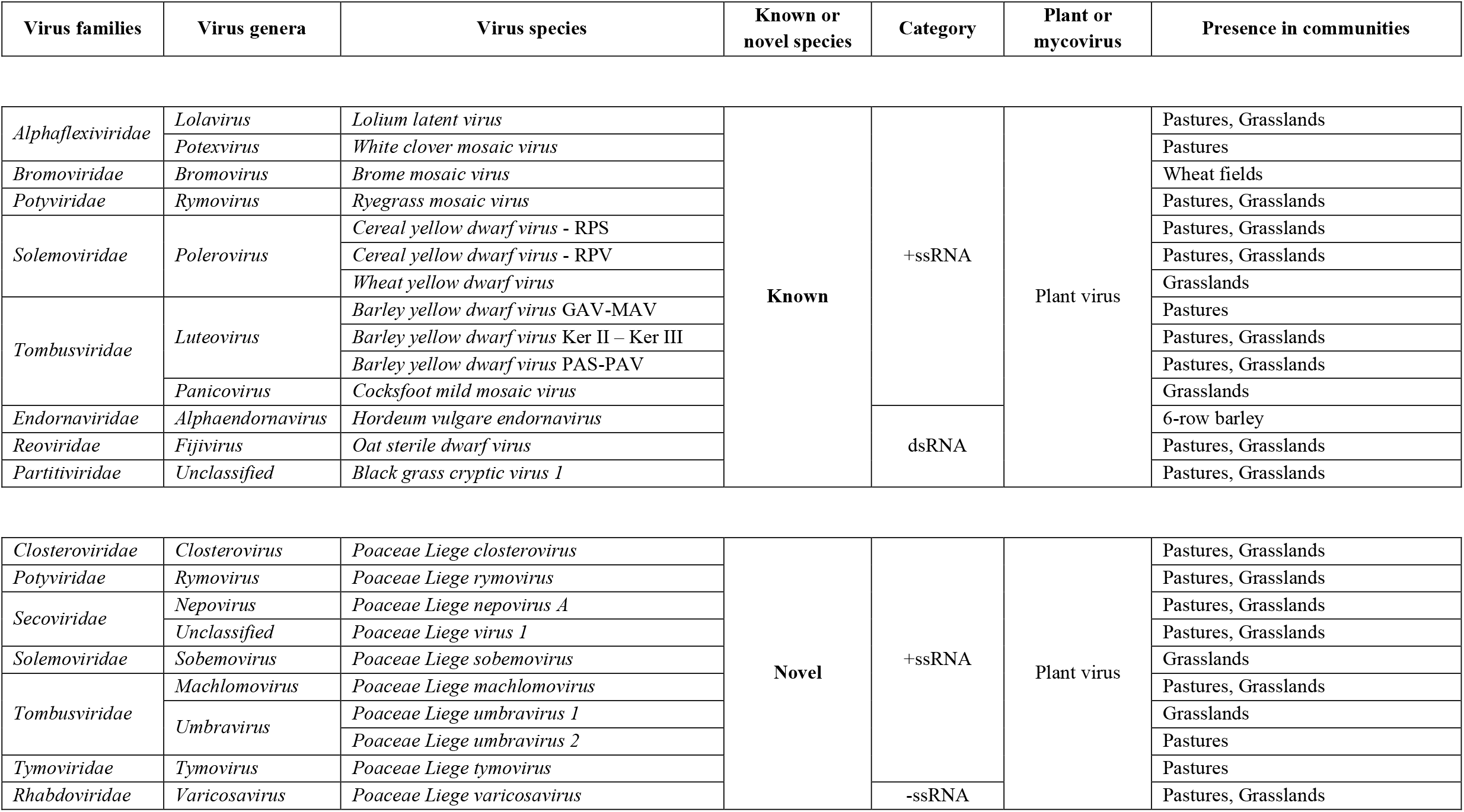
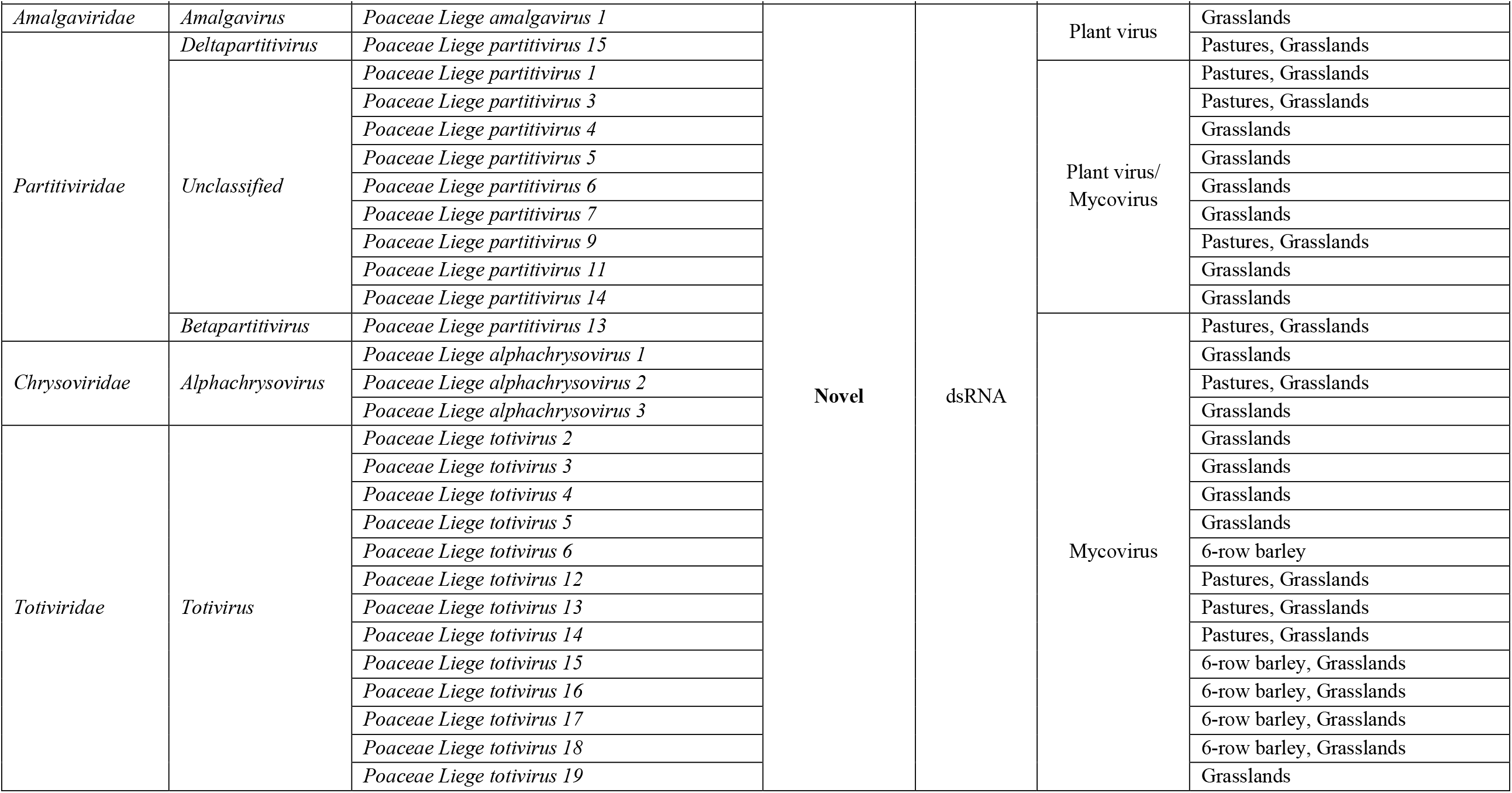
List of the different virus families, genera and species detected in the *Poaceae* communities (cereal fields, grazed pastures, mowed grasslands). For the BYDV species complex, recombination events did not allow to go down to the species levels, and in some cases the species were thus regrouped as GAV-MAV, KerII-KerIII and PAS-PAV.

**Figure 1.**
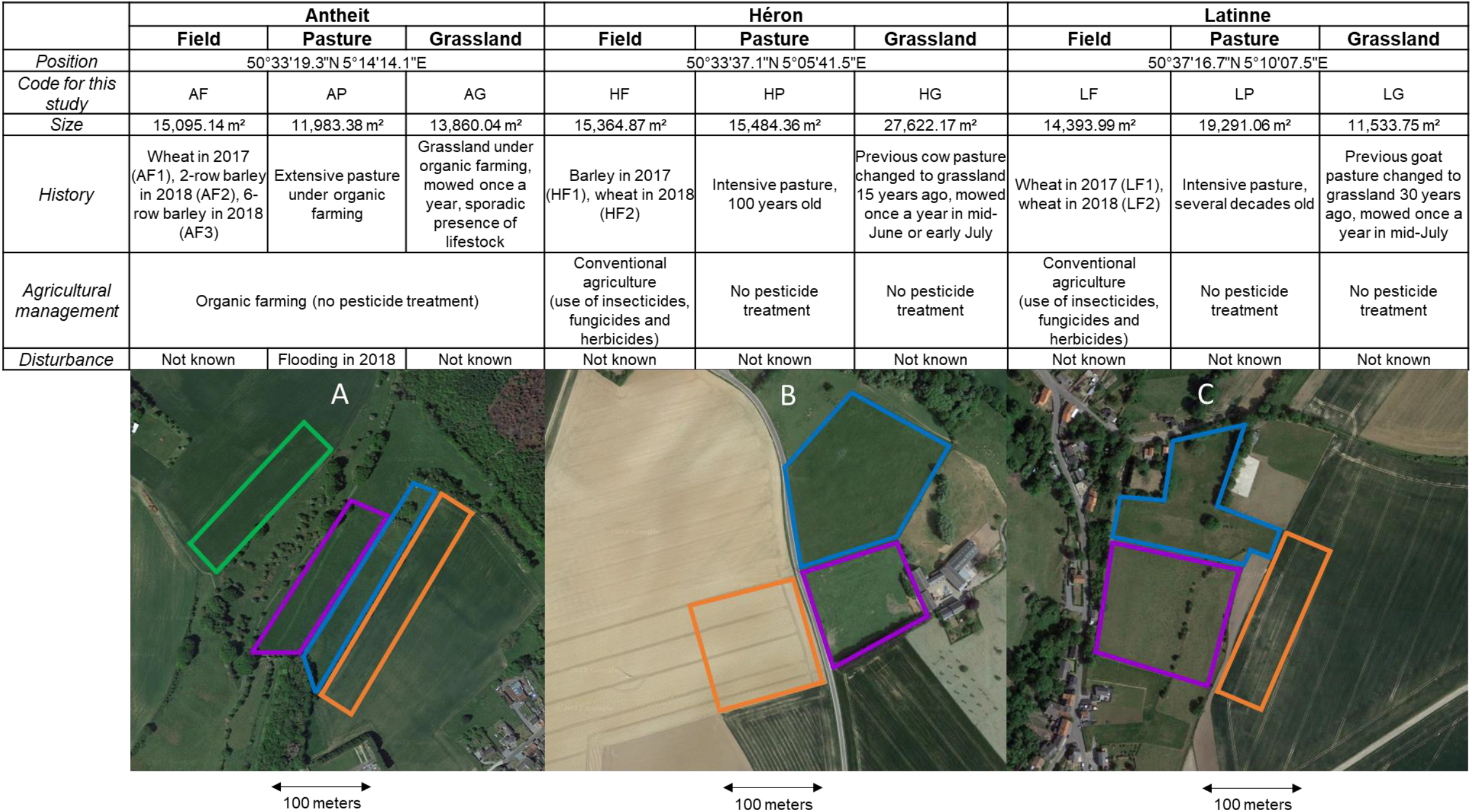
Information table (GPS position, code, size, history, agricultural management, and eventual disturbance) in the different communities and locations studied. Below, a representation of Antheit (A), Héron (B) and Latinne (C) locations, with the three adjacent ecosystems examined: a barley or wheat field (in orange), an intensive pasture (in purple) and a grassland with high biological value (in blue). An additional 6-row barley field (in green) was sampled in 2018 in Antheit.

### Relationship between Poaceae communities and virome richness or composition

Preliminary HTS results revealed contrasted virome richness among plant communities. Significant differences virus taxa richness were observed between cereals fields, pastures and grasslands (p-value < 0.001 at virus family, genus and species levels for the Kruskall-Wallis tests). Very few or no virus species were found in cereal fields, while a diversified virome was observed in more diverse communities with up to 22 virus species detected in grasslands (**Figure 2**). The replicate of Héron grassland in 2017 (code HG1) had a different mowing management and presented the lowest grassland virome (8 virus species). Interestingly, pastures and grasslands did not present significant differences in virome richness at family and genus levels (p-values 0.456 and 0.419, Mann Whitney tests), but the analysis at species level highlighted a more diversified virome in grasslands than pastures (Mann Whitney test, p-value 0.078 but decreased to 0.010 when excluding HG1). Linear regression models were used to show the relationship between plant species richness and virus family richness (virus families = 0.60 + 0.62 grass species, R-sq = 74.3%, p-value 0.000), virus genera richness (virus genera = 0.55 + 0.82 grass species, R-sq = 72.9%, p-value 0.000) and virus species richness (virus species = 0.22 + 1.57 grass species, R-Sq 76.0%, p-value 0.000; see **Figure 2**).

**Figure 2.**
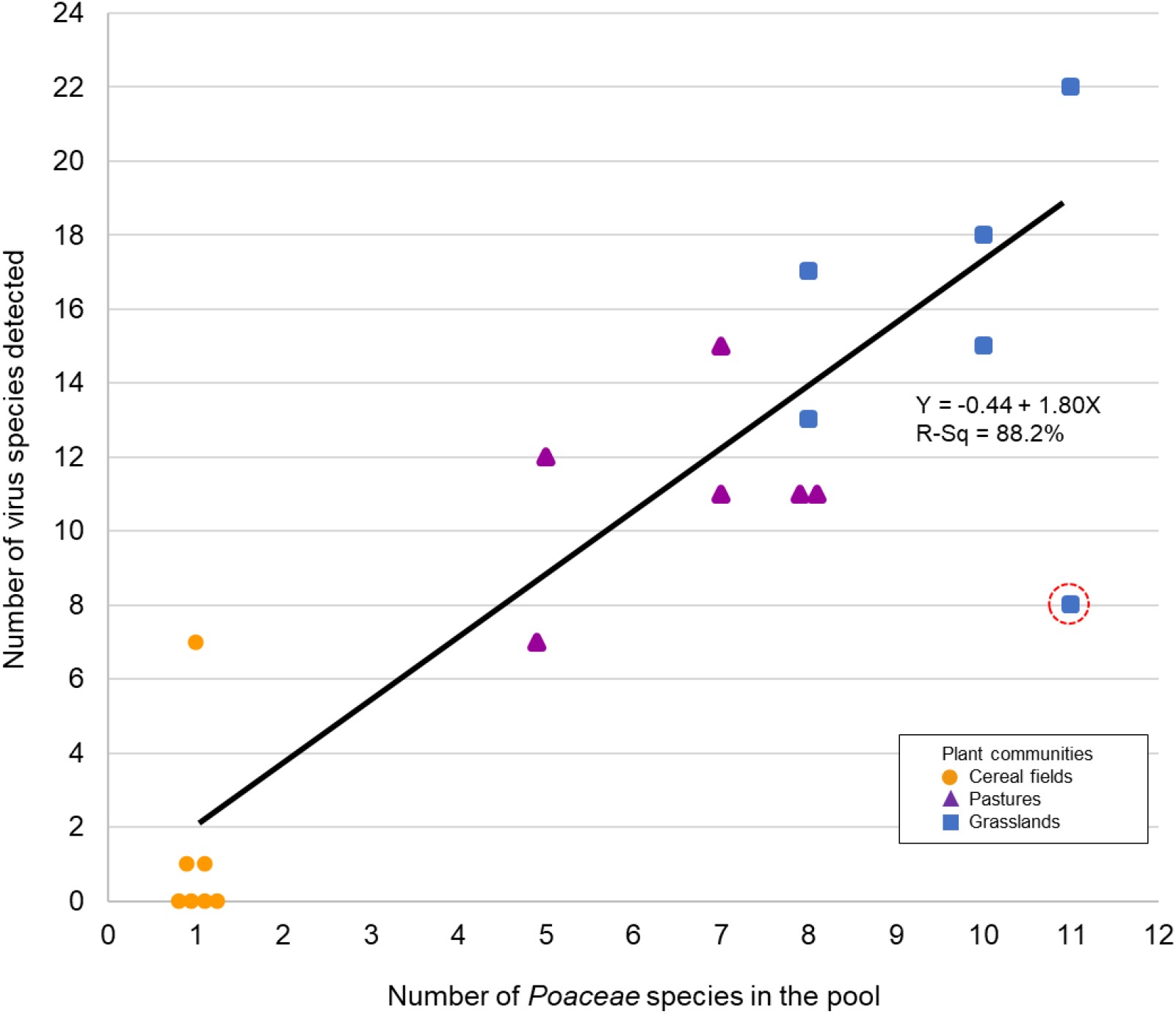
Relationship between *Poacene* species richness and vims species richness among plant communities for the two years studied (cereal fields in orange circle, pastures in purple triangle, grasslands in blue square). The survey area Héron Grassland 2017 with a different mowing management is highlighted with a dotted red circle. TI1e exclusion of Héron Grassland 2017 further improved the R-Sq value, from 76.0% to 88.2%.

Virome composition among *Poaceae* communities were visualized using a network analysis of the virus species identified in each grass community (**Figure 3**), and we performed a clustering analysis in both virus and plant dimensions for the grasslands and pastures to highlight any community clustering or virus co-occurrence patterns (**Figure 4**). Exclusion of cereal fields improved branch length in the clustering model (see **Figure S1** for the three communities clustering) When considering *Poaceae* communities, replicates belonging to the same community but from different sites and years clustered together (**Figures 3 and 4**). We observed two main clusters for the less managed communities: the grasslands from Héron and Latinne, and the pastures to which can be added Antheit grassland (**Figure 4**). The 6-row barley fields clustered with Héron and Latinne grasslands, and the wheat field in Héron formed a single cluster (**Figure S1**) When considering virus species, network analysis highlighted a series of ubiquitous virus taxa detected in several or even all *Poaceae* community types, such as the persistent viruses and mycoviruses (partitiviruses, totiviruses, alphachrysoviruses) and some non-persistent (or acute) viruses (Poaceae Liege nepovirus A, Lolium latent virus, yellow dwarf viruses). In addition to this shared virome, virus species (mostly non persistent viruses) were detected in specific communities, for example brome mosaic virus in wheat crop; ryegrass mosaic virus in pastures; cocksfoot mottle virus in grasslands. Interestingly, the network analysis showed association between *Poaceae* communities and virus biology. Pasture viromes were rather composed by non-persistent viruses (i.e. 66% of the non-persistent virus edges in pastures), while grasslands were more associated with persistent viruses and mycoviruses (i.e. 73% of the persistent virus edges in grasslands) (**Figure 3**). This trend was confirmed by the clustering analysis that structured the virome into two main clusters of co-occurrent viruses mainly associated to each *Poaceae* community (**Figure 4**).

**Figure 3.**
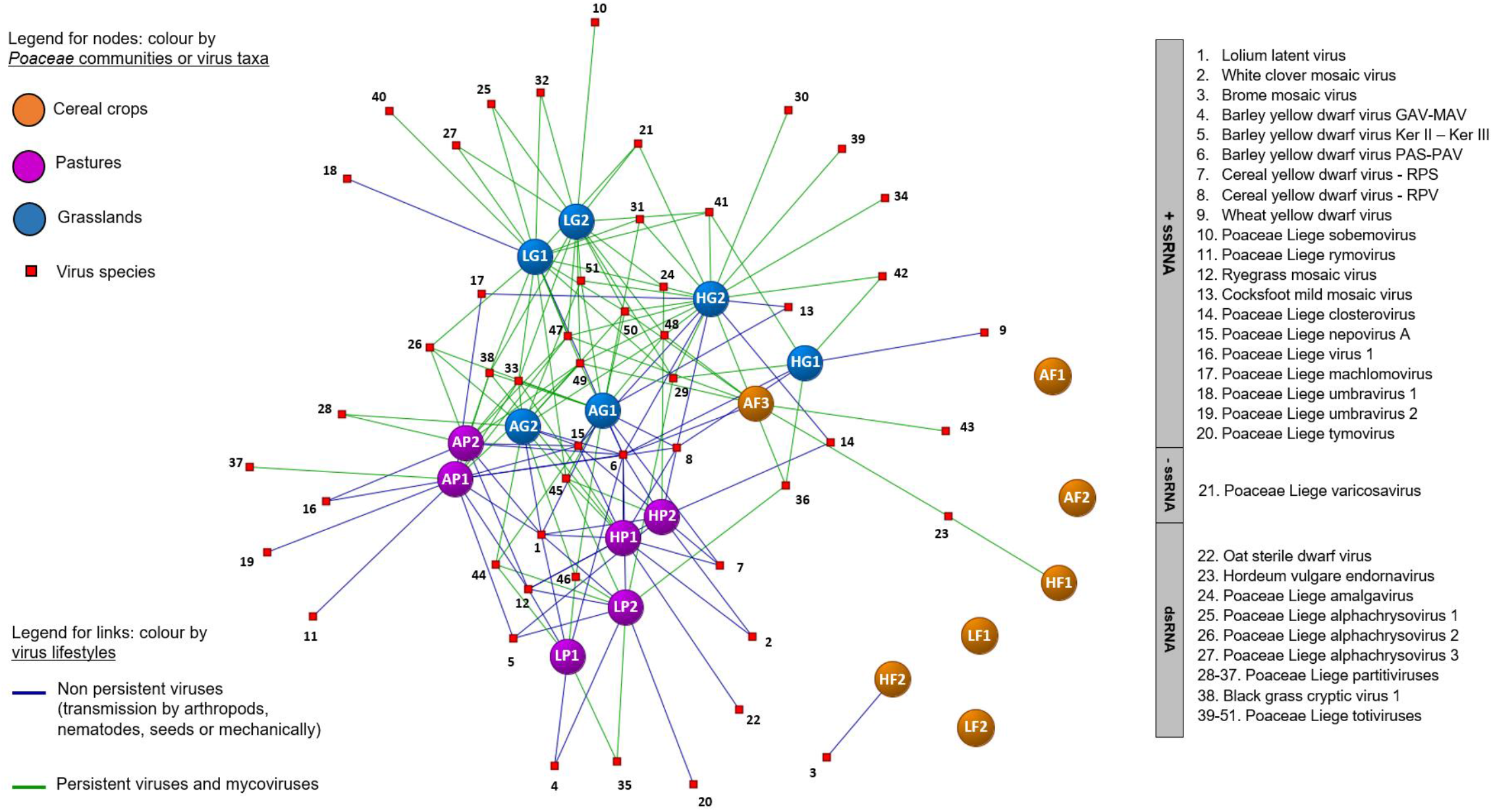
Network anlalysis of the virus species identified in *Poaceae* communities (cereal crops, pastures and grasslands) across the three sites and the two years studied. Nodes correspond to virus species (red square) or plant community (coloured balls). Links are coloured according to the putative lifestyle for each virus species (non-persistent viruses, persistent viruses and mycoviruses).

**Figure 4.**
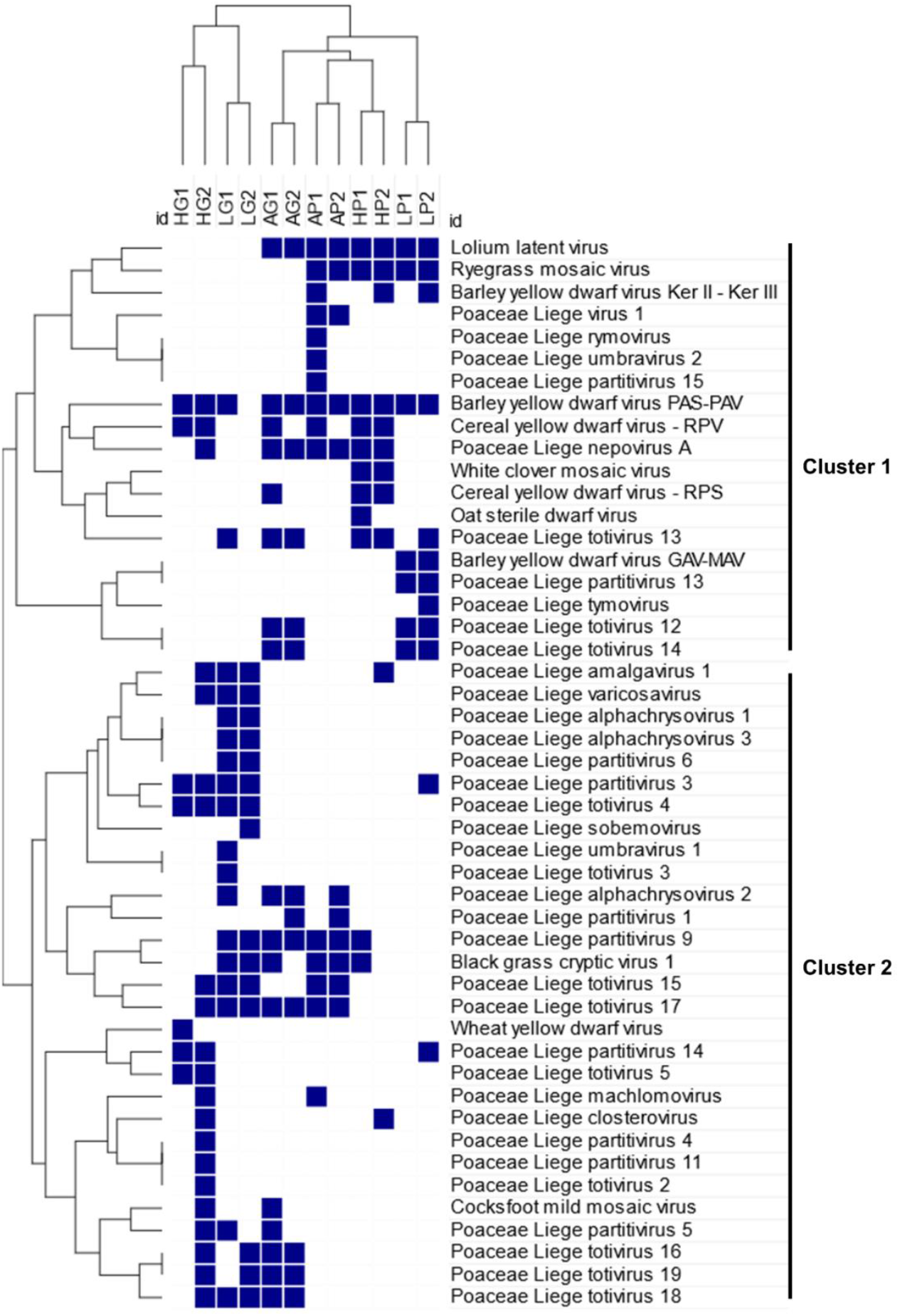
Hierarchical clustering analysis in both plant and virus dimensions. Columns referred to the two different wild *Poaceae* communities (Pastures (P) and Grasslands (G)) examined among sites (Antheit (A), Héron (H) and Latinne (L)) for the two years (2017 (*1*) and 2018 (*2*)). Rows corresponded to the different vims species detected. Two main clusters of co-occurent virus species were highlighted on the right of the dendrogram.

### Higher viral prevalence and multiple infection in less managed plant communities

To complete the first ecological information provided by virome composition in plant pools, we analyzed virus prevalence, co-infection and distribution within the plots. For that, we performed virus detection in individual plants from the Antheit site in 2018 for three viruses detected by HTS: barley yellow dwarf virus-PAV (BYDV-PAV, found in all communities) and two novel secovirids (Poaceae Liege nepovirus A - PoLNVA and Poaceae Liege virus 1 - PoLV1) recently characterized (31) and detected in both pastures and grasslands. Individual plants from community pools (grassland, pasture and 6-row barley field) and from two dominant species (*Lolium perenne, Poa trivialis*) were analyzed. Infirming our hypotheses, we observed high virus prevalence associated with the absence of visible symptoms in less managed communities (i.e. pastures and grasslands). The PoLNVA nepovirus presented high prevalence in both pastures and grasslands, as well as in *L. perenne* and *P. trivialis* (maximum of 88% found in *L. perenne* in pasture). Conversely, we determined contrasted prevalence for BYDV-PAV and PoLV1, depending on the type of plant community or on the plant species (**Table 2**). In the barley field, the hypothesis of high prevalence was not fulfilled, with only 6% of BYDV-PAV prevalence observed. The hypothesis of high co-infection rates in the wild compartment was confirmed, in particular for co-infection by BYDV-PAV and PoLNVA (up to 50% of plants co-infected in the grassland pool). Interestingly, multiple infections by the three viruses were globally detected at a low rate in the samples (0-7%), except for the *P. trivialis* with 16% and 26% of multiple infected plants in grassland and pasture respectively (**Table 2**). These numbers are higher than the predicted co-infection rate (*i*.*e*. the product of individual rates for the three viruses, which gives predictions of 7% for grassland (0.24*0.74*0.4) and 15% for pasture (0.56*0.74*0.36). Concerning single infections, low rates were observed for BYDV-PAV and PoLV1 (0-16%), while single infection by PoLNVA was more prevalent and even reached 68% in *L. perenne* in pasture (**Table 2**).

**Table 2.** A) Virus prevalence of BYDV-PAV and the two novel species (PoLNVA and PoLV1) observed among plant communities and species at the Antheit sites in 2018. Plant community pools were designated as “Global”. Plant species pools were also examined, for *Lolium perenne* and *Poa trivialis*. B) Percentages of plants in single and multiple infection with BYDV-PAV, PoLNVA and PoLV1 in uncultivated plant communities (grassland and pastures, named “Global”) and two wild *Poaceae* species (*Lolium perenne* and *Poa tivialis*), in Antheit in 2018.

| A. Prevalence |  | Percentage of infected plants |  |  |
| --- | --- | --- | --- | --- |
| Plant community | Category | BYDV-PAV | PoLNVA | PoLV1 |
| Field | Global | 6% | 0% | 0% |
| Pasture | Global | 26% | 54% | 4% |
|  | *Lolium perenne* | 6% | 88% | 20% |
|  | *Poa trivialis* | 56% | 74% | 36% |
| Grassland | Global | 72% | 68% | 22% |
|  | *Lolium perenne* | 18% | 84% | 10% |
|  | *Poa trivialis* | 24% | 74% | 40% |

| B. Mixed infection |  | Percentage of infected plants |  |  |  |  |
| --- | --- | --- | --- | --- | --- | --- |
| Type of virus infection | Global Grassland | Global Pasture | *Lolium* Grassland | *Lolium* Pasture | *Poa* Grassland | *Poa* Pasture |
| Single infection BYDV PAV | 16% | 4% | 2% | 0% | 0% | 8% |
| Single infection PoLNVA | 4% | 32% | 58% | 68% | 40% | 22% |
| Single infection PoLV1 | 2% | 2% | 0% | 6% | 6% | 0% |
| Co-infection BYDV PAV - PoLNVA | 50% | 20% | 16% | 6% | 4% | 20% |
| Co-infection BYDV PAV - PoLV1 | 6% | 0% | 0% | 0% | 4% | 2% |
| Co-infection PoLNVA-PoLV1 | 8% | 0% | 10% | 14% | 14% | 8% |
| Infection 3 viruses | 6% | 2% | 0% | 0% | 16% | 26% |
| No infection by any of the 3 viruses | 14% | 40% | 14% | 8% | 16% | 14% |

Virus spatial patterns were analyzed for *P. trivialis* and *L. perenne* in grassland and pasture. The nepovirus (putatively transmitted by seeds, pollen and/or nematodes) was distributed throughout the plots, with a few clusters of non-infected plants (I_a_ = 1.72-1.90, P_a_= 0.02 or 0.03), except for the *P. trivialis* pasture with a random distribution of non-infected spots (I_a_ = 1.00, P_a_= 0.38). On the other hand, viruses transmitted by insects (*i*.*e*. aphids for BYDV-PAV or putatively aphids or leafhoppers for PoLV1) presented some clustering of infected plants (**Figure 5**). SADIE analyses revealed trends of virus aggregation within the plots, in particular for PoLV1 in grasslands for both *P. trivialis* and *L. perenne* (I_a_ = 1.842 and 1.839 respectively, P_a_= 0.0171). Virus distributions were also compared to each other, and contrasted associations were shown between insect-transmitted viruses according to the plant species: mostly positive association for *P. trivialis* (highest positive value of X = 0.54 with p-value = 0.00 for BYDV-PAV and PoLV1 in pasture) and negative association for *L. perenne* (highest negative value of X = -0.62 with p-value = 0.99 for BYDV-PAV and PoLV1 in grassland) (see **Table S5**).

**Figure 5.**
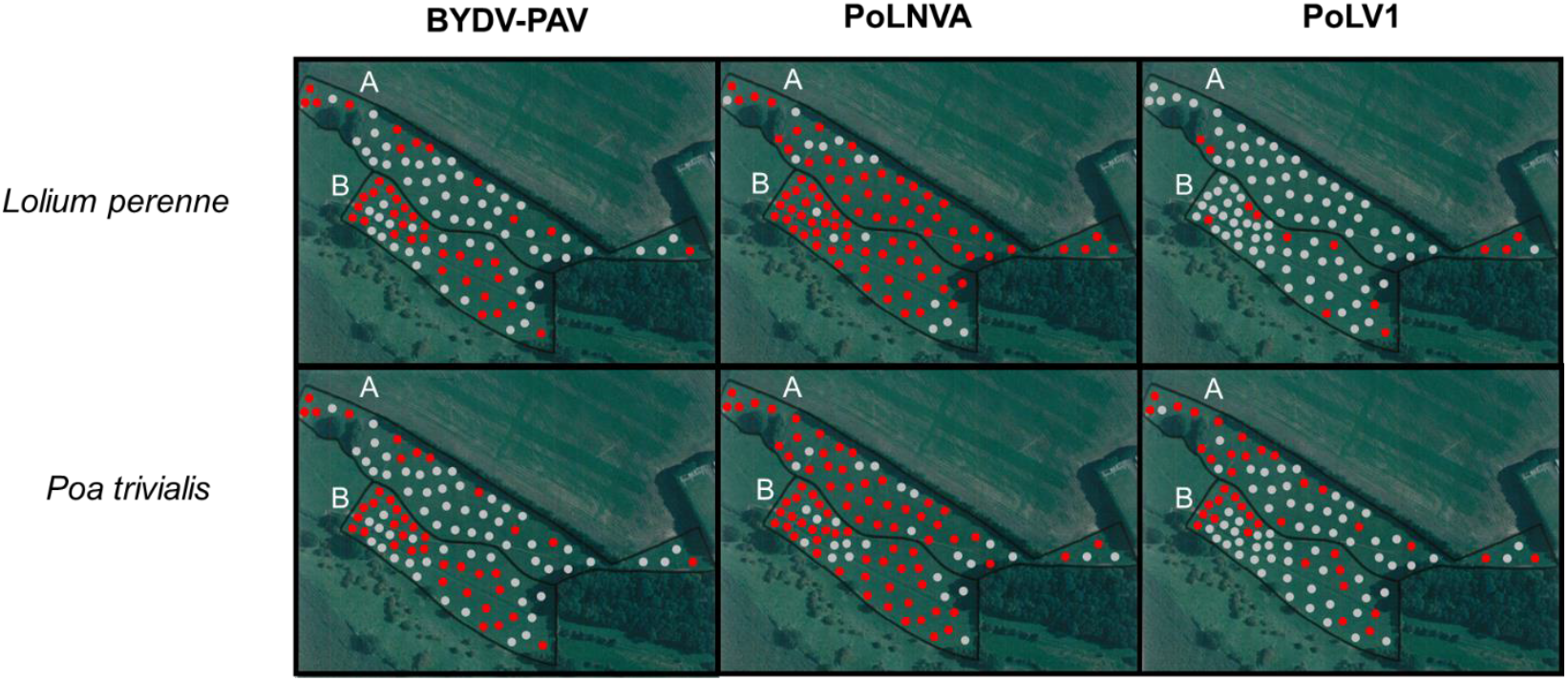
Geographical distribution of infected plants (for BYDV-PAV, PoLNVA and PoLVl) belonging to *Lolium perenne* and *Poa trivialis* in Antheit grassland (A) and pasture (B). Grey and red dots correspond to non-infected and infected plants respectively.

### Large viral genetic structure in wild grasses

Inter-annual comparison of viral sequences within the same plant communities revealed a high genome-level genetic stability for numerous virus species. For instance, 99.4% and 99.8% nucleotide identities were observed for ryegrass mosaic virus (RMV) and for PLNA respectively in Antheit pasture between both sampling years. However, more variability was found for Lolium latent virus (LLV; Genus *Lolavirus*, Family *Alphaflexiviridae*), which made it interesting to examine the virus genetic structure among communities and sites. This analysis was performed in *L. perenne* as this species seemed to be the major host for LLV (higher prevalence and more complete genome obtained as compared to other grass species). Preliminary results of BLASTn searches revealed a high level of variability of LLV contigs that were sharing 70%-99% nucleotide identity with LLV isolates stored in the NCBI GenBank database. Calculating the pairwise identity shared by the RdRp region of LLV isolates detected in this study further confirmed their genetic diversity. Hence, the LLV RdRp pairwise identity matrix (**Table S6**) presented a series of clusters and subclusters with 75-99% nucleotide identity between LLV RdRp sequences recovered from this study. Phylogenetic analyses performed on LLV RdRp sequences (**Figure 6**) showed three main clusters of LLV. Their composition showed various interesting features with closely related RdRp sequences detected: i) in the same plot for the two years studied (*e*.*g*. AP1-C1 and AP2-C5, HG1-C1 and HG2-C1); ii) in pasture and grassland from same location (e.g. AG1-C3 and AP1-C4, HP2-C2 and HG2-C2); (iii) in different sites (e.g. AG1-C2, HP2-C1 and LP2-C2); or iv) detected only once for a given plot and/or location (e.g. HP1-C1 or LP1-C1). In addition, very different RdRP sequences were observed at the same time in the same plot (e.g. AG1-C1, AG1-C2 and AG1-C3), some of which closely related to sequences identified in other sites (e.g. AG1-C2, HP2-C1 and LP2-C2).

**Figure 6.**
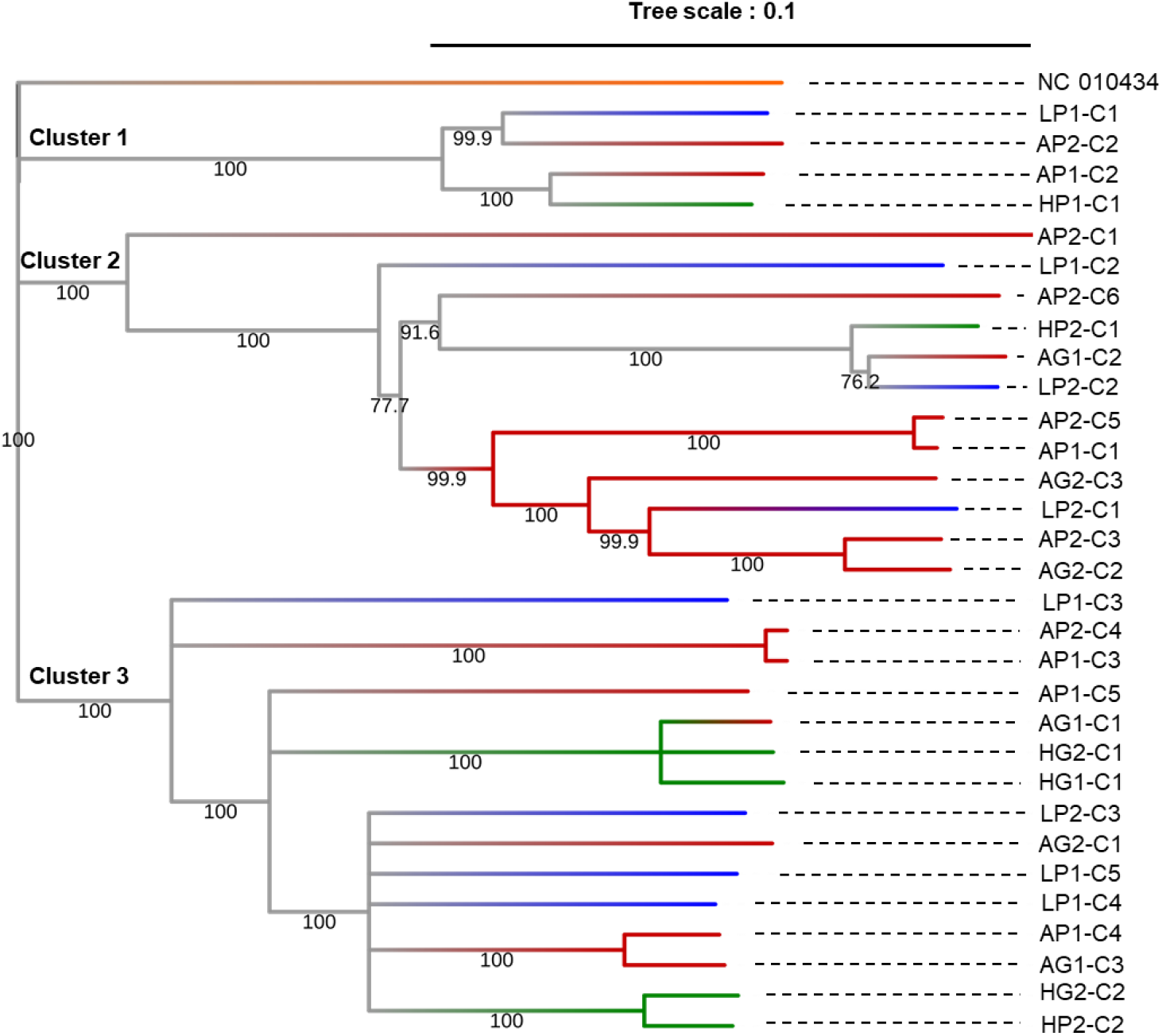
Phylogenetic trees (Neighbor-Joining method, outgroup NC_010434, 1000 boostraps, threshold of 70% occurrence) of the RdRp region of lolimu latent vims in *L. perenne* in different sites (Antheit *(A)* in red Héron *(H)* in green and Latinne (*L*) in blue) and communities (grassland (*G*) and pasture (*P*)) for the two years (2017 *(1)* and 2018 (*2*)) on consensus sequences (contigs de novo assembled *-Cx)*. Samples are compared to the NCBI reference of LLV (isolate Usl, NC_010434) in orange.

## Discussion

The present study has led to significant advances in the identification of the *Poaceae* virome and the host-parasite richness relationships in agro-ecological landscapes. The results have revealed the complex structure of viral communities in nature by combining different ecological analyses (virus richness, network, prevalence, co-infection, spatial distribution and genetic structure). So far, plant virus metagenomics-based studies have focused their ecological analyses on family or genus taxonomic ranks, or examined viral OTUs clustered at a standard level of 90% identity (20, 29, 32, 33). Improving the taxonomic resolution of such studies toward the species level is a crucial point to fine tune ecological analyses such as the estimation of virus(es) genetic structure and diversity (richness and evenness), or even to characterize novel virus species (22). In this study, we strove to obtain as complete virus genomes as possible, covering for many of them all known ORFs described in the corresponding genera or families. The virome could thus be analyzed at species level, allowing more-in-depth ecological analyses (correlation between plant and virus species richness, virus network, phylogeny, etc.). One limitation was for segmented viruses (e.g., partitiviruses). It was not possible to associate the different segments in samples pools containing several partitiviruses and we thus focused the analysis on segment 1 (encoding the RdRp). Noteworthy, hundreds of small viral contigs (*i*.*e*. 300-500 nt) presenting homologies to members of *Chrysoviridae, Endornaviridae, Partitiviridae, Rhabdoviridae* and *Totiviridae* families and to unclassified mycoviruses were also detected in wild *Poaceae*, but could not be classified at the species level. These contigs suggest the presence of additional virus species at low abundance or low titer within the communities.

A diversified virome was thus identified in the Belgian *Poaceae* communities, with at least fifty-one viruses belonging to twenty-one genera and sixteen families (**Table 1**). As the VANA approach can potentially detect both RNA and DNA viruses, the virome of temperate *Poaceae* species appeared in our study to be vastly dominated by RNA viruses. In addition, this *Poaceae* virome was found to be more than two thirds composed of unknown species, in particular persistent and fungal-associated virus taxa from *Chrysoviridae, Partitiviridae and Totiviridae* families. These genera were found in all less managed communities and in one cereal field, highlighting their large distribution in plants and confirming other recent virome-based studies (20, 29, 33, 34).

In this study, three *Poaceae*-based communities with contrasted plant richness due to anthropic management were compared revealing that virome richness increases with the number of grass species within the sampled community. The positive correlation observed between plant and virus species richness agrees with other studies analyzing host and parasite richness in plants and animals (35–37). Globally, our findings are innovative for plant viruses as the few disease ecology studies examining plant pathogens have mainly analyzed foliar fungal pathogens or the relationships between host and virus richness only for some targeted plant viruses (e.g. five viruses in (27), eleven viruses in (28)). A recent Geometagenomics study performed in France and South Africa highlighted that some cultivated areas (that had lower host diversity) could present a greater virus families diversity than uncultivated ones (20). In the present study, comparison of the virus family or genus diversity between communities showed less clear tendencies than using virus species, as illustrated in grasslands versus pastures, making the results difficult to compare with those of (20).

Network and clustering analyses also highlighted different virus lifestyles among the wild *Poaceae* communities. More acute/non-persistent virus taxa were identified in pastures and in Antheit grassland, while Héron and Latinne grasslands harbored more persistent viruses and mycoviruses (**Figures 3 and 4**). Plant diversity and land use can thus influence both the virome richness and composition, as illustrated here by the mowing management. In Héron grassland, mowing date variation changed the virus richness and composition observed in 2017 and 2018, and in Antheit the sporadic presence of livestock in the grassland led to a virus composition closer to pastures. Bernardo et al. (2018) reported that some virus families were significantly associated with cultivated or with unmanaged areas in France and South Africa. Spatial distribution of host populations is expected to be a key determinant of disease dynamics, with an infection risk tending to increase with increasing host population size and connectivity to other populations (38). Infection spread of vector-transmitted viruses might be favoured in dense networks of host populations (39), which can explain why we observed more non-persistent viruses in pastures. By contrast, more diverse, long-term and less connected plant populations as found in grasslands could lead to increase the distribution of persistent viruses that are transmitted vertically via gametes. In cereal fields, the high host density could lead to high virus spread but it may be counterbalanced using pesticides and the yearly sowing of new certified seeds. In particular, the impact of insecticides on the vectors of phytoviruses, and of fungicides on the spatial prevalence of fungi and therefore of mycoviruses are expected to affect the virome, as suggested by the results of (34). This could explain the differences in the virome observed between the organic (in Antheit) and conventional (in Héron and Latinne) cereal fields.

Temporal dynamics in the *Poaceae* virome were also examined and revealed a relative stability with 85% of virus species detected in both years in the permanent plant communities, i.e. grazed pastures and mowed grasslands. This confirmed the absence of significant differences in plant virus population composition observed for a few targeted viruses in natural plant communities (33). Identification of plant viruses down to species levels also allowed the analysis of the intraspecific variability, as explored here with LLV. Several situations were observed, with different virus genomes coexisting in the same studied plant communities, some of them detected both years while others identified a single year only, as well as one situation of closely related genomes detected in different sites (**Figure 6**). Results would suggest potential circulation of virus isolates among communities within the same site, but not frequent exchange among sites. In addition, 18/31 of the LLV variants were not found from one year to another, indicating that the variant diversity was incompletely sampled and is likely larger than reported. This observation is relevant because it shows that the sampling strategy was well adapted for studying the composition at species level but that studying the population genetic structure of a virus would require an additional sampling effort.

Individual plants examination showed contrasted prevalence, co-infections and spatial distributions among plant communities, plant species and virus species. Nematodes-transmitted and single stranded RNA seed/pollen-transmitted viruses such as nepoviruses have been demonstrated to have large host range breath (HRB) (40), in line with the observation that numerous *Poaceae* species were found infected by the novel nepovirus (31). Moreover, the two other viruses studied, BYDV-PAV and PoLV1, presented more contrasted prevalence depending on plant community or species (**Table 2**). Examination of transmission modes can, at least partially, contribute to explain these differences. Unlike nepoviruses, BYDV-PAV is transmitted by aphids (and novel PoLV1 is close to waikaviruses that are also insect-vectored). Complex interactions are involved in the insect transmission of plant viruses, in particular vector feeding behavior and attraction/deterrence of vectors for host plants (41). These effects are expected to be even more complex in natural compartments composed of multiple plant and insect species.

In this study, grasslands and pastures presented different plant richnesses, with a series of plant species specifically growing in grasslands and not in pastures, and that represented potential hosts for insects. Combined to other factors such as vector feeding preference for some plant species, presence of more dicots diversity in grasslands than pastures, or different mowing management, this can contribute to higher prevalence in grassland than pasture. In addition, various interspecific interactions among insect vectors (competition, co-existence, etc.) differing according to the plant species can potentially impact virus distribution within the plot (**Figure 5**), as illustrated by SADIE analyses (e.g. negative association between BYDV-PAV and PoLV1 in *Lolium perenne*). The over-representation of mixed infection compared to the predicted co-infection rate, as observed in *Poa trivialis*, suggested that the viruses are not circulating independently and some factors such as co-transmission, assistance, or attraction of vectors could have an effect. Results in individual plants, all asymptomatic, confirmed that virus infection is very frequent in nature and generally unapparent (as illustrated and reviewed in (42–44)). BYDV-PAV prevalence was higher in uncultivated areas compared to the cropping system, while previous virome studies (20, 27) and ecological hypotheses predicted an increased pathogen prevalence with increase in host abundance (e.g. in cultivated areas) (45, 46). This lower prevalence could reflect successful management of BYDV-PAV in the cereal crops (e.g., insect vector control, planting period). Conversely, pastures and grasslands are mostly composed by perennial grasses that can harbored viruses regardless of vector pressure of a given year.

Besides its implications in an ecological perspective, the identification of novel plant viruses down to species levels also contributed to discuss and relativize the species demarcation criteria proposed by ICTV, in particular for persistent viruses and mycoviruses (*Alphachrysovirus, Partitivirus, Totivirus, Victorivirus* genera). Indeed, ICTV genome-based species demarcation criteria are very contrasted between taxa: from 80-90% amino acid sequence identity for partitiviruses to 53-70% for alphachrysoviruses, 60% for victoriviruses and even 50% for totiviruses. Biological criteria such as the host species are also considered (47). In the present study, most of the novel virus species identified presented low identity levels (14-80% amino acid identity for RdRp and CP regions, see **Table S4**) compared to known species in the same genus/family. Their status as novel species leaves no doubts for partitiviruses as they were far below the demarcation criteria, but novel totiviruses were more complex to discriminate as they presented identity levels (14-62% aa identity) sometimes close to the demarcation threshold. Before the advent of metagenomics, the ICTV criterion of 50% amino acid identity was sufficient because the limited number of known species were sufficiently distinct (e.g. only 30% identity between saccharomyces cerevisiae virus L-A and saccharomyces cerevisiae virus L-BC in the 717 aa region of highest similarity) (47). However, the numerous *Totivirus* species identified here and in other metagenomics studies (34, 48) presented less clear tendencies that could eventually suggest to reconsider demarcation criteria for the genus (e.g. harmonizing rules with victoriviruses that also belong to the *Totiviridae* family and present a demarcation threshold of 60% aa identity).

In summary, this large-scale study revealed a diversified and largely unknown virome in cultivated and non-cultivated plant communities, with high viral prevalences and co-infection observed in less managed communities, and the virome richness increasing with the grass species richness in the plant communities. Virome network highlighted different virus lifestyles among wild *Poaceae* communities, illustrated the influence of plant diversity and land use on the composition of viral communities. The examination of vectors (arthropods, nematodes, fungi, etc.) is a crucial element to include in virus metagenomics studies to analyze spill-over and spill-back between wild and agricultural reservoirs. In addition, plant genotyping of different species may provide information on virus adaptation to various hosts (as explored here with LLV). The approach developed here for *Poaceae* can be extended to other plant families (e.g. *Solanaceae, Fabaceae*) in order to provide a global overview of the virus communities present in these mixed species communities, and to progress in understanding the ecology of plant viruses across agro-ecological interfaces, a domain still in its infancy.

## Material and Methods

### Selection of the Poaceae communities and sites

We characterized the virome of three co-occurring *Poaceae-*dominated communities that represent a gradient of plant species richness and management intensity: (i) cereal crops (low plant species richness, intense management); (ii) pastures grazed throughout the year by cattle (moderate species richness, moderate management), and (iii) naturalistic grasslands managed only by annual mowing (high species richness, low management). We studied these community types (**Figure 1**) across two years (2017 and 2018) at three replicated sites ten kilometers apart (Antheit, Héron, Latinne) within the National Park “Burdinale-Mehaigne” (Province of Liège, Belgium), where climate is temperate. The cereal crops included two-row and six-row barley (*Hordeum vulgare* subsp. *vulgare* L. and subsp. *hexastichum* (L.) Celak, respectively) and winter wheat (*Triticum aestivum* L.). The crop fields were cultivated conventionally (with commercial fertilizers and pesticides) at Héron and Latinne and with organic practices (organic fertilization and no pesticides) in Antheit. In contrast, the grazed pastures were sown a century ago and contain four dominant grass species, all of which are cool-season C3 perennials native to Eurasia: *Agrostis capillaris* L., *Dactylis glomerata* L., *Lolium perenne* L. and *Poa trivialis* L. Lastly, the grasslands were communities with high conservation value, containing up to fourteen *Poaceae* species and additional forbs. These grasslands had been mowed once a year in mid-July for the past fifteen--thirty years (see **Table S1**), except in Héron which was mowed in mid-May in 2017.

### Sampling strategy

To capture the virome of these cool-season *Poaceae* communities, we surveyed them twice--from mid-May to mid-June of 2017 and 2018 (18 surveys: 3 community types x 3 replicates x 2 years). In 2018, we sampled an additional six-row barley field (AF-3) adjacent to the pasture plot in Antheit because it presented interesting symptoms of fungal infection. Thus, there were 19 surveys in total. For each survey, we randomly placed 50 quadrats (1m x 1m) in the target area (1.1 – 2.7 ha). Within each quadrat, we inventoried all *Poaceae* species and noted any viral or fungal symptoms. We harvested one randomly chosen individual (one stem with leaves) of each species identified within that specific quadrat. Harvested plant material was kept cool on ice freeze packs in the field and then stored at -80°C later that day. To compare the virome captured in each survey, we prepared 19 corresponding pools of 50 individual plants each, based on botanical inventory of the 50 quadrats visited in each survey (see **Table S2**). Each pool reflected the relative abundance of the *Poaceae* species encountered in that survey. For instance, 20% of the grasses within the 2017 Antheit Grassland survey were *L. perenne* (47/232 plants sampled, see **Table S1**), so this species was represented in the 50-sample pool by 10 plants. Non-*Poaceae* species were not surveyed.

### Purification of virus-like particles and nucleic acid extraction

Both DNA and RNA viruses were targeted in this study. The 19 pools of plants were prepared by virus-like particles (VLPs) enrichment, followed by the extraction and sequencing of virion-associated nucleic acids (VANA) as described previously in (31). To obtain the VLPs, 200 mg of frozen tissues from each individual plant were pooled into a filtered bag (*i*.*e*. 10 g for fifty plants) and then grinded in 70 ml of Hanks’ buffered salt solution (HBSS, composed of 0.137 M NaCl, 5.4 mM KCl, 0.25 mM Na_2_HPO_4_, 0.07 g glucose, 0.44 mM KH_2_PO_4_, 1.3 mM CaCl_2_, 1.0 mM MgSO_4_, 4.2 mM NaHCO_3_), using a tissue homogenizer. The homogenised plant extracts were centrifuged at 3,200 g for 5 min. Supernatants were further centrifuged at 8,200 g for 3 min. Resulting supernatants were then filtered through a 0.45 μm sterile syringe filter and 25 ml of supernatant were put into an ultracentrifuge tube. Then, a sucrose cushion, made of 3 ml of 30% sucrose in 0.2M potassium phosphate pH 7.0, was added. Extracts were then centrifuged at 148,000 g for 2.5 hours at 4°C using the SW 28 Ti rotor (Beckman). Resulting pellet was resuspended overnight at 4°C in 1.5 ml of HBSS. Unencapsidated nucleic acids were eliminated by adding 15 U of bovine pancreas DNase I (Euromedex) and 1.9 U of bovine pancreas RNase A (Euromedex, France) to 200 µl of resuspension. Samples were incubated at 37°C for 90 min. VANA were then extracted from 200 μl of resuspended virions using a PureLink Viral RNA/DNA Mini Kit (ThermoFischer Scientific, Belgium) following the manufacturer’s protocol.

### Library preparation and high-throughput sequencing

Reverse transcription, Klenow fragment treatment, and amplification of the viral DNA and RNA using barcoded PCR primers were then prepared according to the protocol of (49). Amplification products were cleaned up using the Nucleospin Gel and PCR clean up (Macherey-Nagel, Germany). Ten libraries with unique multiplex identifier linkers (MID, described in (50)) were further assembled and sequenced at GIGA-Liège University (Belgium) using the NEBNext Ultra II DNA library prep kit (New England BioLabs) and then sequenced on the Illumina NextSeq 500 platform with 2 × 150 nt reads with a target of 10 million sequences per pool of ten libraries (i.e. 1 million sequences per 50-plant pool).

To control proper extraction of RNA and DNA viruses (and to deal with cross-samples contaminations, a series of blank (water) and alien controls were used in each batch of sequenced samples. An alien control, which have been recommended in recent guidelines (51), corresponds to a plant sample infected by an virus (called alien virus) that cannot infect the studied plant samples. For this study, potato leaf roll virus (genus *Polerovirus, RNA virus*) and banana streak virus (genus *Badnavirus*, DNA virus) were selected. The presence of reads from alien viruses in the samples correspond to cross-contamination between samples and can support the establishment of a thresholds below which presence of a virus is considered as background cross-contamination.

### Sequence analyses and annotation

After quality check, raw reads were demultiplexed and cleaned according to their multiplex identifier (MID) linkers with Cutadapt (52). Then, reads were *de novo* assembled into contigs (SPAdes assembler, 2.5% mismatches allowed, minimum size of 35nt) (53) on the Durandal cluster (ULiège, Belgium). Contigs were annotated using BLASTn and BLASTx (54) and NCBI nucleotide (nt) and protein (nr) databases respectively. Viral BLASTx hits (conservative e-value < e^-20^ cut-off) were isolated for further analyses carried out on Geneious Prime 2019.2.1 (https://www.geneious.com). In order to obtain complete or nearly complete virus genomes, we assembled the virus contigs and then mapped the reads on these contigs (Geneious mapper, 5 iterations, medium-low sensitivity, 5% mismatches and 5% gaps allowed). Novel virus species were further tentatively identified following host and genomic ICTV species demarcation criteria for each virus family (host range and nucleotide/protein identity percentage on RdRp and coat protein). In order to obtain robust results, we performed the analyses (virome richness, network, phylogeny) on complete (or nearly complete) virus genomes, excluding the small contigs (less than 500 nt) that did not allow to form complete virus sequences.

### Viral genetic structure analysis

We examined the population genetic structure of Lolium latent virus (LLV) over years, communities and sites. The abundance of this virus in almost all the pools including *Lolium* samples (e.g. up to 80% of total reads in Antheit grassland in year 2) allowed to analyze LLV intraspecific genetic structure, phylogeny and variants among and within plant communities. As the complete genome of LLV was not obtained for all samples, the analysis focused on the RNA-dependant RNA polymerase (RdRp) gene that was better covered (corresponding to nt 88-5,277 on the NCBI reference genome, NC_010434). After obtaining de novo LLV contigs for each sample, a multiple nucleotide alignment of the 31 contigs was performed on MUSCLE (55, 56) with 8 iterations, and a Nigh tree was built on Geneious (Neighbor-Joining tree, Tamura-Nei model, outgroup NC_010434, 1,000 bootstraps and a support threshold of 70% occurrence in the consensus tree).

### Statistical and network analyses

Viromes at family, genus and species level were inventoried from the 19 pools. Statistical analyses were performed on Minitab19 to determine whether viromes richnesses (*i*.*e*. number of virus taxa identified in each plant community) were significantly different between plant communities and anthropogenic managements. Population structure was analyzed, through the distribution normality and variance homogeneity. As the two hypotheses were not fulfilled for all the conditions, we used the Kruskal-Wallis test to analyze all communities, and the Mann-Whitney test to compare two community categories (e.g., pastures and grasslands). Both tests were performed with a 95% confidence level. In addition, network analyses of the virome composition (i.e. the different virus taxa constituting the virome) in the different *Poaceae* communities were implemented on ORA-LITE software (netanomics.com). The aim of this analysis was to visualize how virus species are distributed among the communities, and to determine any association between virus taxa and plants, e.g. comparing persistent and non-persistent (acute) virus lifestyles, according to the definition of (57) (see **Table S7** for the virus-community matrix used). The same matrix was used to perform hierarchical clustering analyses on the Morpheus software (Pearson correlation analysis, default parameters, https://software.broadinstitute.org/morpheus), recursively merging objects (*i*.*e. Poaceae* communities and virus species) based on their pairwise distance.

### Virus prevalence, co-infection and spatial distribution in individual plants

Total RNA was extracted according to (58). Reverse transcription was carried out with Tetro RT enzyme (Bioline). The 20 µl RT reaction mix consisted of 2.5 µl of total RNA (at 300 ng/µl), 10 µl of RNAse-free water, 4 µl of 5x RT buffer, 1 µl of random hexamers (50 µM), 1 µl of dNTP mix (10 mM), 0.5 µl of RNAse OUT (40U/µl) and 1 µL of Tetro enzyme (200U/µl). The reactions were then incubated as follows: 25°C for 10 min, 45°C for 30 min, 85°C for 5 min and then placed directly on ice. Amplification of the cDNA was performed using Mango Taq enzyme kit (Bioline) in 25 µl reaction mix composed of 2.5 µl of cDNA, 12.5 µl of RNAse-free water, 5 µl of 5x PCR buffer, 0.5 µl of dNTP mix (10 mM), 1 µl of each primer (20µM), and 0.5 µL of Mango Taq enzyme (5U/µl). Primers (see **Table S8**) were previously designed for each targeted virus (BYDV-PAV, novel PoLNVA and novel PoLV1) from the HTS data using Geneious Prime 2019.2.1 Software (https://www.geneious.com). Thermal cycling corresponded to: 94°C for 4 min, 35 cycles at 94°C for 45 sec, Ta for 1 min and 72°C for 30 sec, with a final 72°C extension for 10 min. PCR products were analyzed by electrophoresis on a 1% agarose gel in TAE buffer stained with GelRed® Nucleic Acid Gel Stain (Biotium) and visualized under UV light.

### Spatial Analysis by Distance IndicEs (SADIE)

Spatial distribution was analyzed for BYDV, PoLNVA and PoLV1 in *L. perenne* and *P. trivialis* through the SADIE (Spatial Analysis by Distance IndicEs) technique using SADIEShell Version 2.0 and N_AShell Version 1.0. (Rothamsted Experimental Station, Harpenden Herts, United Kingdom). This approach is used to study spatial patterns in count-based data where location are known and test their statistical significance (59). Comparing observed patterns of samples with two extremes (crowding and regularity), an index of aggregation (I_a_) was calculated to quantify the presence and degree of clustering. Values of I_a_ = 1 indicate a random spatial distribution, I_a_ > 1 reveals aggregation of counts into clusters and I_a_ < 1 points out a regular or uniform spatial distribution (60, 61). Statistical significance of aggregation was evaluated by the null hypothesis (P_a_) that the counts are arranged randomly with respect to one another (significant if P_a_ < 0.05 or P_a_ > 0.95 meaning aggregative or regular pattern respectively) (62). Clustering indices can be used to determine an association index X in order to compare different viral populations within same location and evaluate if they occur close together in similar habitats (plant community x plant species), or conversely if they are segregated from one another. This analysis is associated to a significance test under the null hypothesis of no association (with p < 0.025 or p > 0.975 for a significant positive or negative association respectively) (63). For BYDV-PAV and PoLV1, we hypothesized aggregation spots related to insect presence in the field.

## Data availability

Raw sequencing reads were submitted in the NCBI Sequence Read Archive (SRA), with the BioProject accession number PRJNA882095. All data related to virus genome assembly and identification, and their NCBI GenBank accession numbers are listed in **Table S4**. All other data are provided in the Supplementary information section of this paper.

## Acknowledgements – Funding

This work was supported by the Fonds de la Recherche Scientifique - FNRS under Grant No 1.1.309.19F06 and J.0149.20. We would thank the Gembloux Plant Pathology Laboratory team for the plant sampling and samples processing.

