## Supplementary material for "Long term anthropic management and associated loss of plant diversity deeply impact virome richness and composition of *Poaceae* communities": Table S8

### Supplementary Table S8: Primers used for targeted virus detection by RT-PCR

Primers were previously designed for each targeted virus (BYDV-PAV, novel PoLNVA and novel PoLV1) from the HTS data using Geneious Prime 2019.2.1 Software (https://www.geneious.com). They are listed with their respective annealing temperatures (Ta) in the table hereunder.

| Primers | Sequence (5’-3’) | 5’ Position | Ta (°C) | Amplicon size |
| --- | --- | --- | --- | --- |
| BYDV-F  BYDV-R | CCCAGTCTATCGCAATGCCCAGC  GGTTCCGGTGTTGAGGAGTCTAC | 3104  3483 | 55°C | 379 bp |
| PoLNVA-F  PoLNVA-R | ACCCTCAAGTTCTTTCCACTT  ACTCCCTCTCCAGTATTGAA | 3775  4150 | 63°C | 375 bp |
| PoLV1-F  PoLV1-R | TGTGTCGGGAAATAAACTACAAGCA  GCAAAAGAGCCAAACTGGAATGGTA | 3251  3607 | 56°C | 356 bp |
