## Supplementary material for "Long term anthropic management and associated loss of plant diversity deeply impact virome richness and composition of *Poaceae* communities": Figure S1

**Supplementary Figure S1**. Hierarchical clustering analysis in both plant and virus dimensions. Columns referred to the different Poaceae communities (Fields (F), Pastures (P) and Grasslands (G)) examined among sites (Antheit (A), Héron (H) and Latinne (L)). Rows corresponded to the different virus species detected.

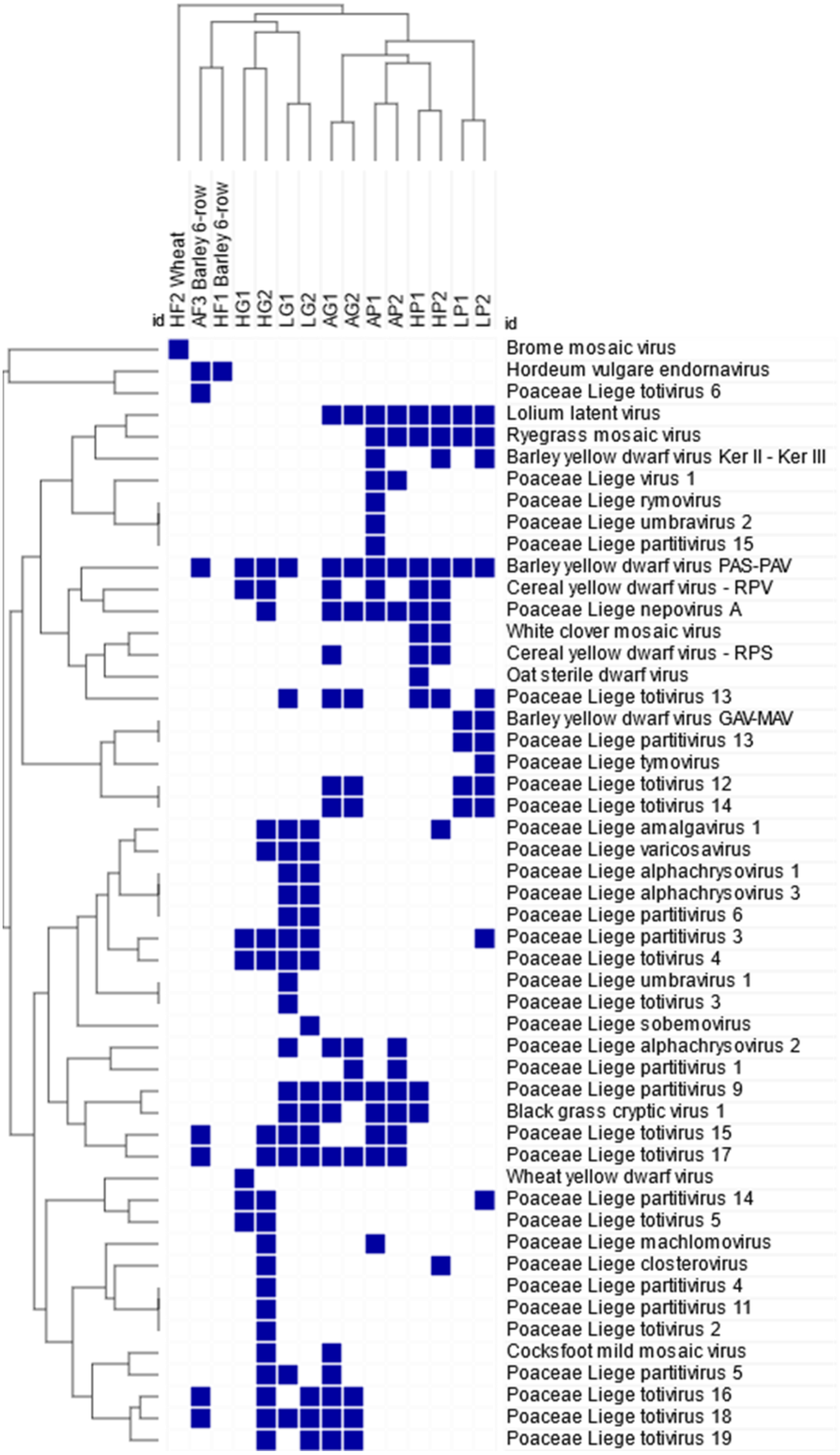
